# In Silico Discovery of Small Molecule Modulators Targeting Ebola Virus VP 35 and VP 40 protein

**DOI:** 10.1101/2025.01.31.636006

**Authors:** Yang Liu, Chen Zhu, Mei chen, Daoyu Guo

**Affiliations:** Department of Internal medicine and Pediatrics, clinical college of Qilu medical university; Department of Cardiology, affiliated hospital of Qilu medical university; Department of Internal medicine and Pediatric, clinical college of Qilu medical university; Department of Nursing science, clinical college of Qilu medical university; Department of Neuroscience, clinical college of Qilu medical university

## Abstract

Ebola virus (EBoV) proteins VP35 and VP40 are acknowledged as significant pharmacological targets for disrupting viral replication, virion assembly, and the budding process from host cells. We employed molecular docking based throughput virtual screening (HTVS), standard precision (SP), and extra precision (XP) docking to discover candidate compounds. Subsequently, molecular dynamics simulations, and ADMET analyses to identify potential small molecular compounds that could inhibit the Ebola virus. Among 210,590 compounds from the SPECS database, we found the compounds AO-022/43452438, AN-465/43369333, and AN-465/43369198 were optimal candidates for the EBOV-VP35 protein, while AN-465/43369160, AN-465/4341136, and AN-465/43369952 also emerged as optimal compounds for the EBOV-VP40 protein. Furthermore, we employed the energy decomposition technique, MM-GB/SA, to identify the top ten amino acids that significantly contribute to the binding interactions. ADMET result implicated that the six compounds have potential druggability, but special attention should be given to AN-465/43369333 due to potential cardiac toxicity. In conclusions, the six compounds and their key interacting amino acid sites identified in this study may serve as a valuable platform for the future development of novel therapeutics targeting the Ebola virus.

## Introduction

Preliminary findings from high-throughput screening (HTS) have yielded varying success rates, identifying hundreds of candidate compounds with potential efficacy against EBOV (1). EBOV proteins VP35 and VP40 have been recently identified as two key multifunctional proteins.VP35 protein facilitates viral replication by perturbing type I interferon signaling, and has NTPase and helicase-like activities (2, 3), while VP40 protein contributes to viral assembly and budding (4). Thus, the majority of druggable targets against EBOV focus on inhibiting these two specific proteins. In our study, we employed an in silico methodology that included molecular docking-based HTVS, molecular dynamics simulations, and pharmacological assessments to identify promising compounds targeting the EBOV proteins VP35 and VP40.

## Results and Discussion

The screening process was conducted in a sequential manner, retaining 10% of the molecules at each stage. In the virtual screening of two proteins, we commenced by filtering the compound database to eliminate high-energy conformations and compounds that did not adhere to the five established rules of drug-like properties, resulting in a total of 131,100 molecules. Following this initial filtration, molecular docking was performed. The HTVS docking phase allowed us to select conformations, reducing the pool to 13,110 molecules. SP docking further narrowed the selection to 1,311 compounds. Ultimately, XP docking obtained 132 molecules (Fig. 1 *A*). We calculated the MM-GBSA binding energies for these molecules. Three candidates with the most favorable MM-GBSA scores are chemical 1 (-51.35kcal/mol), chemical 2 (-47.08kcal/mol), and chemical 3 (-45.04kcal/mol) for the VP35 protein. Meanwhile, the leading three molecules are chemical 4 (-44.93kcal/mol), chemical 5 (-43.47kcal/mol), and chemical 6 (-43.15 kcal/mol) in the case of the VP40 protein. The 2D structure of six candidate molecules were shown in *Fig 1B*.

**Fig 1.**
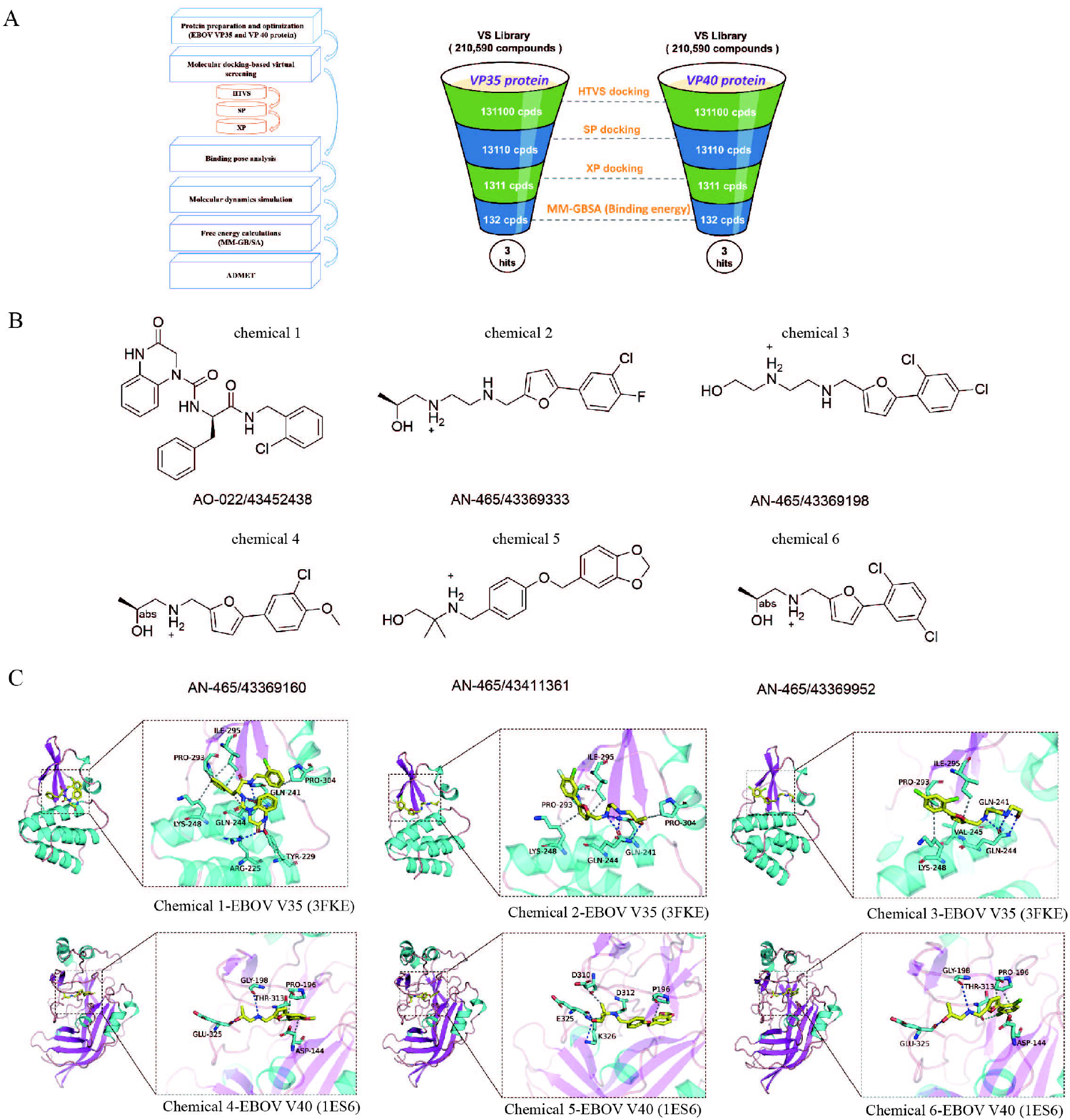
Chemicals identified by the hierarchical virtual screening approach based on molecular docking. (A) Work flowchart of virtual screening. (B) The 2D structure of each chemical. (C) Docking poses of chemicals in the active pocket in EBOV VP 35(PDB ID: 3FKE) and VP40 protein (PDB ID: 1ES6); The left image provides an overall view, while the right image offers a partial view. The yellow sticks represent small molecules, the cartoon depicts proteins, the blue dashed lines indicate hydrogen bonds, the gray dashed lines represent hydrophobic interactions, and the cyan dashed lines illustrate halogen bonds..

Interaction poses, including hydrogen bonds, hydrophobic interactions, and halogen bonds, were calculated among the six chemical-protein complexes (*Fig 1C*). Subsequently, we conducted kinetic simulations on these six specific complexes to enhance predictive accuracy. Root Mean Square Deviation (RMSD) fluctuations of the six complexes, which include chemicals 1 to 3 associated with the VP35 antigen and chemicals 4 to 6 associated with the VP40 antigen, were analyzed throughout the simulation period, along with the fluctuations of their respective ligands. After 60 ns of simulation, the RMSD for five of the systems—chemical 1/2/3-EBOV-VP35, and chemical 4/5-EBOV-VP40 antigen—reaches a stable state, indicating that these complexes are securely bound. In contrast, the chemical 6-EBOV-VP40 antigen complex shows an increasing RMSD in the later stages of the simulation, suggesting poorer convergence and weaker binding stability. The RMSD values for the ligands alone show that the four chemicals 1/3/4/5 display minimal RMSD variations when interacting with the protein. This implies that these small molecules can securely attach within the protein’s active site (Fig. 2*A* and Fig. 2*B*).

**Fig 2.**
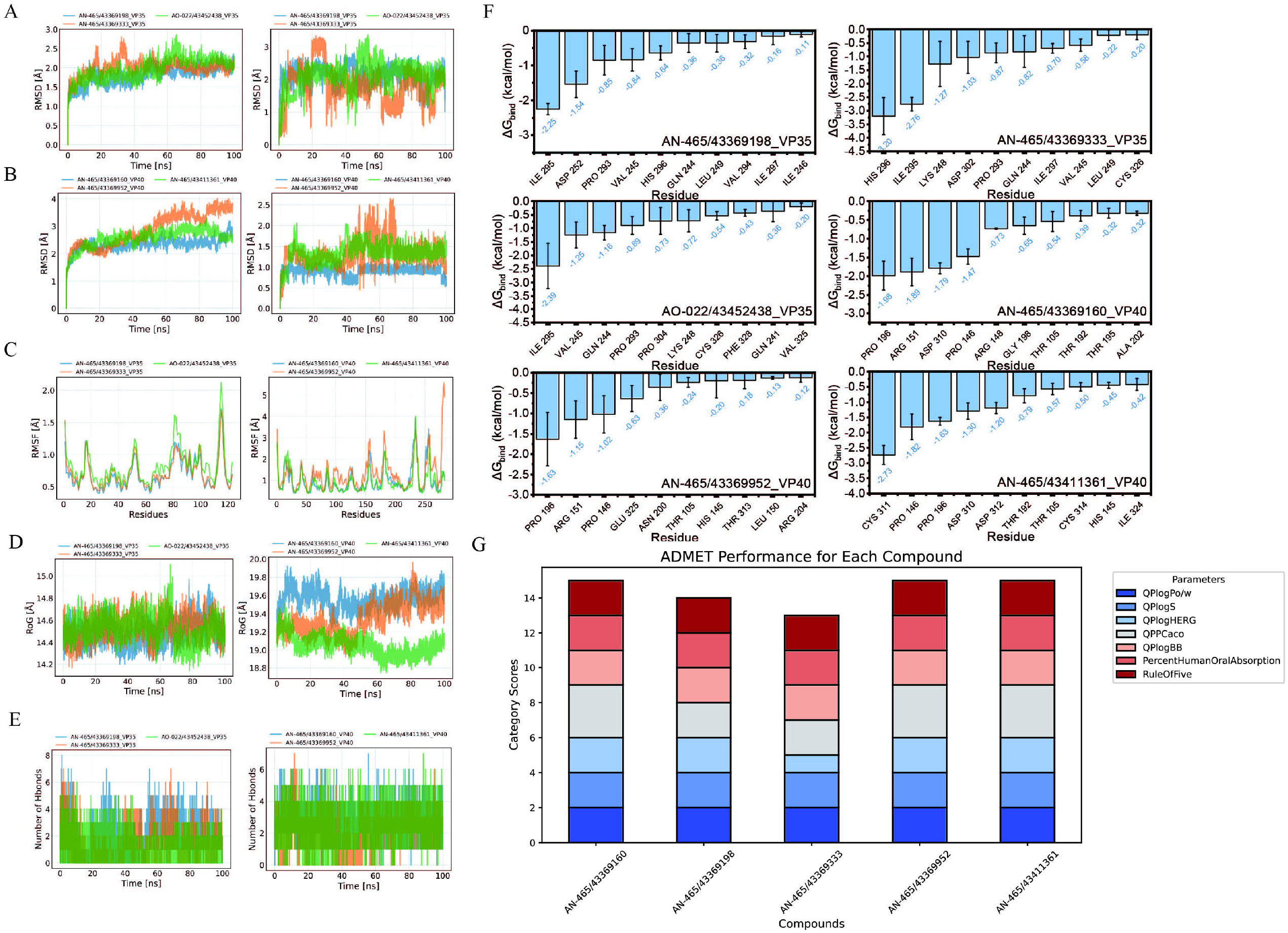
Molecular dynamics simulation and ADMET. (A, B) The root mean square deviation (RMSD) of the complex and the ligand varies over time during the molecular dynamics simulation process. (C) The root mean square fluctuation (RMSF) is calculated based on molecular dynamics simulation trajectories. (D) the variation in the gyration radii of small molecules and protein complexes over time during molecular dynamics simulations. (E) changes in the number of hydrogen bonds between small molecules and proteins during molecular dynamics simulations. (F) the top 10 amino acids contributing to the binding of small molecules and proteins. (G) ADMET.

Generally, the binding of a chemical to a protein results in a decrease in the protein’s flexibility, which in turn enhances stability and effects enzymatic activity. Root Mean Square Fluctuation (RMSF) serves as an indicator of protein flexibility during molecular dynamics simulations. The results represented that, with the exception of a localized region, the RMSF values for the protein remain below 2 Å, indicating a high degree of rigidity in the protein’s overall structure, likely attributable to the interaction with small molecules. Importantly, following the binding of EBOV-VP40 to chemical 6, the observed reduction in RMSF is less pronounced than that seen with the binding of chemical 4 and chemical 5. This suggests that chemical 4/5 exerts a more substantial influence on the protein’s flexibility compared to chemical 6 (Fig. 2*C*).

The gyro-radius serves as an indicator of the compactness of composite systems. The gyro-radii of six complexes exhibited temporal variations. Notably, chemical1/2/3_VP35 demonstrated considerable fluctuations and amplitudes, suggesting that their binding differences are minimal. In the context of the VP40 system, chemical 6_VP40 displayed an upward trend, indicating that the system has not reached stability with the binding of the chemical 6. Conversely, chemical 4/5_VP40 maintained stability throughout the simulation period, reflecting a strong binding affinity between these two systems, particularly for the chemical 5_VP40 complex (Fig. 2*D*).

Hydrogen bonding represents one of the most robust forms of non-covalent interactions. In the context of a 100 ns molecular dynamics simulation, we quantified the number of hydrogen bonds formed between ligand molecules and proteins. The formed hydrogen-bonding interactions include chemical 1_VP35 complex (0 to 2), chemical 2/3-VP35 complex (1 to 2), chemical 4/5-VP40 complex (4 to 5) and chemical 6-VP40 complex (1 to 2) (Fig. 2*E*). These results evidenced that the interaction with EBOV-VP35 is not predominantly facilitated by hydrogen bonding, whereas the interaction with VP40 is primarily mediated through hydrogen bonds. Notably, the complexes chemical 4-VP40 and chemical 5-VP40 complexes exhibit stable binding, with a substantial contribution from hydrogen bonding in these interactions. Furthermore, we employ the MM-GB/SA energy decomposition methodology to identify the ten amino acids that significantly influence the binding affinity between small molecules and the EBOV-P35/40 proteins. These identified amino acids exhibited the greatest contributions to the interaction dynamics between the six candidate small molecule compounds and EBOV-VP35/40 protein (Fig. 2*F*).

The drug-likeness and pharmacokinetic properties of identified chemicals was estimated in terms of absorption, distribution, metabolism, excretion, and toxicity (ADMET) characteristics. All compounds show significant promise for drug development based on essential metrics (Fig. 2*G*). The Log *P* (QPlogPo/w) and solubility (QPlogS) values are within an acceptable range, indicating that these compounds have a balanced level of lipophilicity and hydrophilicity. This balance aids in cell membrane penetration and distribution in water-based environments. Predictions regarding HERG channel blockade (QPlogHERG) suggest that most compounds are within a safe range, although AN-465/43369333 (chemical 2) may present a risk of cardiac toxicity and needs further assessment. The Caco-2 permeability (QPPCaco) data indicate that most compounds have good intestinal absorption, with some showing particularly high permeability. The brain/blood distribution coefficient (QPlogBB) predictions are also within a reasonable range, suggesting that these compounds may have suitable brain permeability, making them appropriate for development as non-central nervous system drugs while reducing the risk of excessive central toxicity. Furthermore, the predicted oral absorption rates exceed 75%, indicating high efficiency for oral administration. All compounds adhere to five Lipinski’s rule, demonstrating favorable drug-like characteristics. Overall, these compounds show excellent potential across key ADMET indicators, but special attention should be given to the HERG value of AN-465/43369333 (Table 1).

**Table 1.**
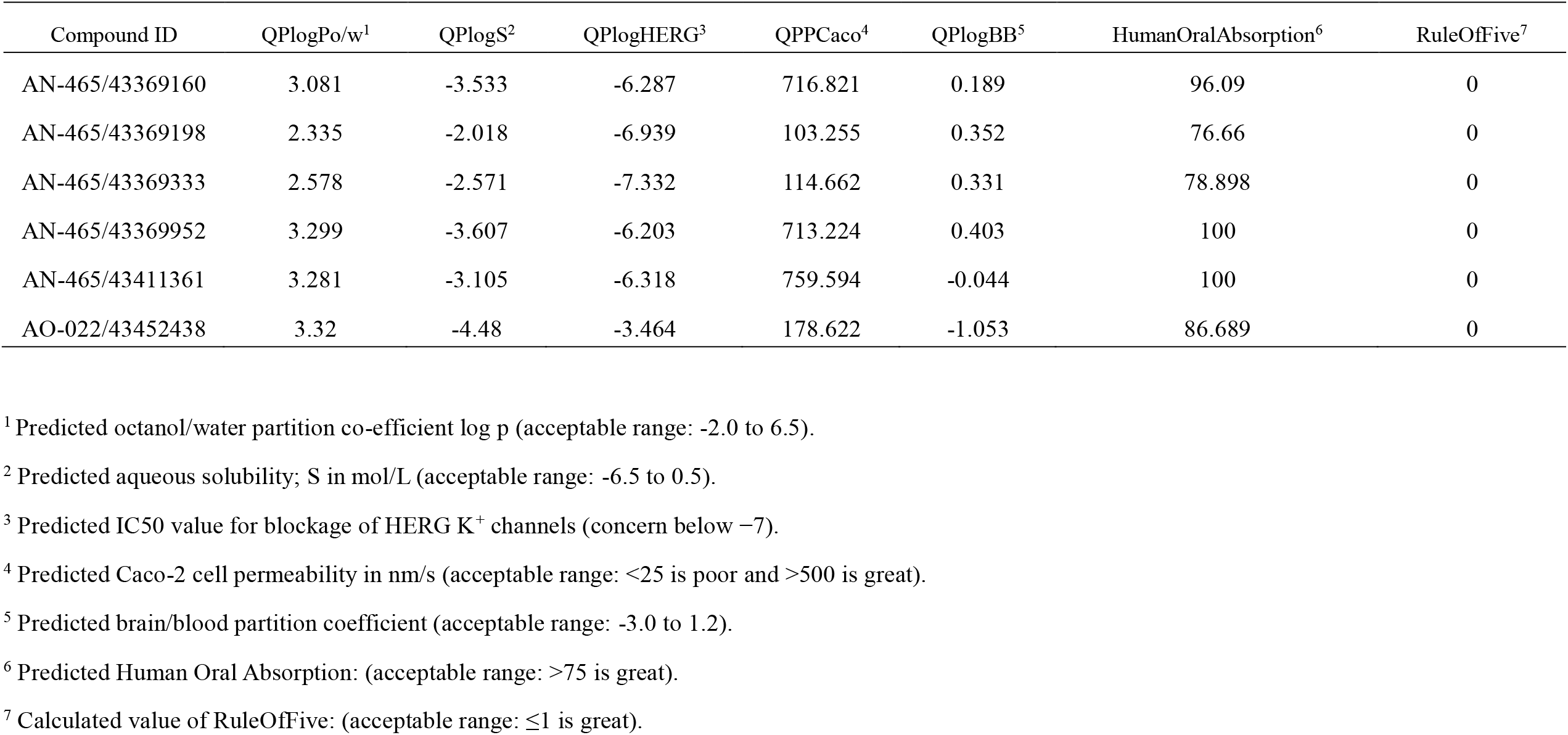
ADME and pharmacological parameters prediction for hits using QikProp.

| Compound ID | QPlogPo/w<sup>1</sup> | QPlogS<sup>2</sup> | QPlogHERG<sup>3</sup> | QPPCaco<sup>4</sup> | QPlogBB<sup>5</sup> | HumanOralAbsorption<sup>6</sup> | RuleOfFive<sup>7</sup> |
| --- | --- | --- | --- | --- | --- | --- | --- |
| AN-465/43369160 | 3.081 | -3.533 | -6.287 | 716.821 | 0.189 | 96.09 | 0 |
| AN-465/43369198 | 2.335 | -2.018 | -6.939 | 103.255 | 0.352 | 76.66 | 0 |
| AN-465/43369333 | 2.578 | -2.571 | -7.332 | 114.662 | 0.331 | 78.898 | 0 |
| AN-465/43369952 | 3.299 | -3.607 | -6.203 | 713.224 | 0.403 | 100 | 0 |
| AN-465/43411361 | 3.281 | -3.105 | -6.318 | 759.594 | -0.044 | 100 | 0 |
| AO-022/43452438 | 3.32 | -4.48 | -3.464 | 178.622 | -1.053 | 86.689 | 0 |

<sup>1</sup> Predicted octanol/water partition co-efficient log p (acceptable range: -2.0 to 6.5).
<sup>2</sup> Predicted aqueous solubility; S in mol/L (acceptable range: -6.5 to 0.5).
<sup>3</sup> Predicted IC50 value for blockage of HERG K<sup>+</sup> channels (concern below -7).
<sup>4</sup> Predicted Caco-2 cell permeability in nm/s (acceptable range: <25 is poor and >500 is great).
<sup>5</sup> Predicted brain/blood partition coefficient (acceptable range: -3.0 to 1.2).
<sup>6</sup> Predicted Human Oral Absorption: (acceptable range: >75 is great).
<sup>7</sup> Calculated value of RuleOfFive: (acceptable range: ≤1 is great).

In conclusions, the six compounds and their key interacting amino acid sites identified in this study may serve as a valuable platform for further development as innovative therapeutic agents.

## Methods

Small molecule compounds used for screening were obtained from the SPECS database. Virtual screening were achieved through the VSW module developed by Schrödinger incorporates three docking algorithms, including HTVS, SP, and XP docking. Molecular dynamics simulations and energy decomposition technique (MM-GB/SA) were employed to estimate the binding affinity and key interacting amino acids between candidate compounds and EBOV VP35/40 proteins. Finally, ADMET characteristics were calculated. Extended methodology is detailed in S*I Appendix*.

## Data, Materials, and Software Availability

Software and database information are detailed in S*I Appendix*. The raw data generated during the current study are available from the corresponding author on a reasonable request.

