## supplemental methods and tables for "In Silico Discovery of Small Molecule Modulators Targeting Ebola Virus VP 35 and VP 40 protein"

### Methodology

#### Methods

#### Crystal structure of EBOV-VP 35/40 protein

Crystal structure of the Ebola VP35 interferon inhibitory domain (3FKE) and the matrix protein VP40 (1ES6) were obtained from RCSB Protein Data Bank (RCSB PDB) (1, 2).

#### Virtual Screening

The SPECS database offers several advantages, including extensive compound information, structural and chemical diversity, potential biological activity, and accessibility. These features provide numerous options and opportunities for drug discovery. In this study, virtual screening was performed on 210,590 compounds from the SPECS database.

#### Database preparation

This study utilized 210,590 small chemical molecules from the SPECS database for virtual screening. This database offers extensive compound information, diverse structures, and a chemical space with potential biological activity, thereby providing numerous options and opportunities for drug discovery.

All molecules underwent a three-dimensional (3D) structural conversion prior to docking, a process completed in the LigPrep module of the Schrödinger software package. This primarily involved energy minimization of all molecules under the OPLS4 force field, utilizing the Epik method for precise protonation prediction and eliminating any potential salt ions present in the molecular library. The purified molecules were then employed for subsequent molecular docking.

#### Protein construction and processing

For the VP35 and VP40 proteins, data were downloaded from the PDB database, with PDB IDs 1ES6 and 3FKE, respectively. Subsequently, the Protein Preparation Wizard module in the Maestro 13.0 software package was utilized to prepare these proteins. This preparation involved adding missing side chain atoms and charges, removing all water molecules, adding hydrogen atoms to the protein system, optimizing the orientation of the hydrogen atoms, accurately predicting the protonation states of the amino acids in an environment with a pH of 7.2, and minimizing the energy of the entire structure using the OPLS4 force field.

**Preparation for the docking box**

The docking box is utilized to restrict the search space for small molecules, typically encapsulating the active site to accurately define the docking process. The docking box file for the active pocket is generated using the Receptor Grid Generation module in Schrödinger 2021-4 software. Based on the prepared protein structure, the SiteMap module is utilized to predict binding sites, with the top-ranked pocket identified as a potential active pocket. The docking box file for the active pocket is generated using the Receptor Grid Generation module in Schrödinger 2021-4 software. The pocket predicted by the sitemap is selected as the center of the box, resulting in a grid file with a volume of $25 \times 25 \times 25$ Å$^3$ (the docking box file).

**Structure-based virtual screening**

The structure-based virtual screening was conducted using the Virtual Screening Workflow (VSW) module in Schrödinger 2021-4 software, which integrates various tools, including QikProp, LigPrep, Glide, and MM/GBSA. As ligand preparation and molecular property calculations were completed prior to pharmacophore screening, this process primarily employed the Glide and MM/GBSA methods.

Molecules obtained through pharmacophore filtering were directly utilized for structure-based virtual screening. The receptor grid generation module was employed to create grid files for docking. During the docking process, the conformations of the compounds must meet the requirement of forming hydrogen bonds with at least two of the three hydrogen bond donors or acceptors located on the hinge of the receptor protein (JAK3 kinase protein): GLU-903:O, LEU-905:O, and LEU-905:H.

The docking process utilized three levels of precision algorithms in Glide: HTVS, SP, and XP, each offering increasing levels of accuracy. All three algorithms were employed in this procedure, following a similar docking protocol that progressively enhanced precision to identify the top 10% of compounds based on their scores for the subsequent round of docking. Ultimately, the XP algorithm was configured to output three conformations for each docking, which were then used for MMGBSA calculations. The MM/GBSA method applied in virtual screening was a non-MD sampling technique, representing a static calculation.

**Molecule dynamics stimulation**

The molecular dynamics simulation was conducted using AMBER 22 software (3). Prior to the simulation, the BCC charges of the small molecules were calculated with the antechamber module (4, 5). Subsequently, the small molecules and proteins were characterized using the GAFF2 small molecule force field and the ff14SB protein force field, respectively (6, 7). The LEaP module was employed to add hydrogen atoms to each system, and the systems were solvated in an octahedral TIP3P solvent box, with a periodic boundary set to 10 Å (8). Sodium ($Na^+$) and chloride ($Cl^-$) ions were added to neutralize the system's charge, and finally, the topology and parameter files for the simulation were generated.

Molecular dynamics simulations were conducted using AMBER 22 software (3). Prior to the simulation, the system underwent energy minimization, which involved 2,500 steps of the steepest descent method followed by 2,500 steps of the conjugate gradient method. Subsequently, the system was heated for 200 ps at constant volume and a controlled heating rate, gradually increasing the temperature from 0 K to 298.15 K. To maintain the temperature at 298.15 K, a 500 ps NVT (constant number of particles, volume, and temperature) ensemble simulation was performed to ensure a uniform distribution of solvent molecules within the solvent box. Finally, a 500 ps equilibrium simulation was conducted under NPT (constant number of particles, pressure, and temperature) conditions. The composite system was then subjected to a 100 ns NPT ensemble simulation under periodic boundary conditions. During the simulation, the non-bonded cutoff distance was set to 10 Å. The Particle Mesh Ewald (PME) method was employed to calculate long-range electrostatic interactions (9), while the SHAKE method was used to constrain the bond lengths of hydrogen atoms (10). Additionally, the Langevin algorithm was implemented for temperature control (11), with a collision frequency (γ) set to 2 ps^-1. The system pressure was maintained at 1 atm, the integration time step was 2 fs, and trajectories were saved every 10 ps for subsequent analysis.

**Free energy calculations (MM-GB/SA).**

The binding free energy between proteins and ligands in all systems is calculated using the MM/GBSA method (12, 13). In this study, molecular dynamics (MD) trajectories ranging from 90 to 100 nanoseconds are utilized for the calculations, and the specific formula is as follows:

$$\Delta G_{bind} = \Delta G_{complex} - (\Delta G_{receptor} + \Delta G_{ligand}) \quad (1)$$

$$= \Delta E_{internal} + \Delta E_{VDW} + \Delta E_{elec} + \Delta G_{GB} + \Delta G_{SA}$$

In formula (3), $\Delta E_{internal}$ represents internal energy, $\Delta E_{VDW}$ indicates van der Waals interactions, and $\Delta E_{elec}$ denotes electrostatic interactions. The internal energy comprises bond energy ($E_{bond}$), angle energy ($E_{angle}$), and torsional energy ($E_{torsion}$); $\Delta G_{GB}$ and $\Delta G_{GA}$ are collectively referred to as solvation free energy. Among these, $G_{GB}$ is the free energy of polar solvation, while $G_{SA}$ is the free energy of non-polar solvation. For $\Delta G_{GB}$, the GB model developed by Nguyen et al. (15) is employed for calculations (igb = 2). The non-polar solvation free energy ($\Delta G_{SA}$) is calculated based on the product of surface tension (γ) and solvent-accessible surface area (SA), with $\Delta G_{SA} = 0.0072 \times \Delta SASA$ (16). The change in entropy is neglected in this study due to high computational resource consumption and low accuracy (12).

**ADMET predictions**

ADMET predictions are conducted using the QikProp module in Schrödinger 2021-4 software and ADMETlab 3.0. The QikProp module provides predictions for QPlogPo/w, QPlogS, QPlogHERG, QPPCaco, QPlogBB, and human oral absorption.

**The software and hardware used for virtual screening**

Protein processing tools: Schrödinger 2021-4 software (Protein Preparation Wizard module); Preprocessing the compound library: Schrödinger 2021-4 software (LigPrep module); Molecular docking simulation: Schrödinger 2021-4 software (Glide module); Free energy calculation (MM-GB/SA): Schrödinger 2021-4 software (Phase module); Visualization software: PyMOL 2.5 (Academic licence); Computer hardware resources: AMAX Server, 56 cores CPU*2: Intel Xeon Platinum 8280; Chemical molecular database: SPECS database (Approximately 210,590 compounds are included)

**Supplemental Table 1:** Screening Results of the VP35 Protein Based on Docking

| Name | TPSA | docking score | MMGBSA dG Bind | Molecular weight | Hydrogen bond donors | Hydrogen bond acceptors | Number of rotatable bonds | AlogP |
|---|---|---|---|---|---|---|---|---|
| AO-022/43452438 | 90.54 | -4.772 | -51.35 | 462.9401 | 3 | 3 | 6 | 4.1209 |
| AN-465/43369333 | 57.43 | -5.015 | -47.08 | 327.8094 | 2 | 2 | 8 | 1.349 |
| AN-465/43369198 | 57.43 | -5.222 | -45.04 | 330.2369 | 2 | 2 | 8 | 1.4304 |
| AP-124/41669777 | 74.49 | -5.551 | -44.68 | 317.4116 | 3 | 3 | 6 | 0.86 |
| AN-465/43369600 | 57.43 | -5.188 | -44.55 | 354.2749 | 2 | 2 | 8 | 1.3362 |
| AO-476/43380647 | 139.51 | -4.734 | -44.4 | 383.4535 | 2 | 4 | 5 | 3.9432 |
| AQ-390/42708960 | 141.65 | -4.931 | -44.22 | 380.4223 | 3 | 5 | 5 | 2.4777 |
| AN-329/41402659 | 124.6 | -4.816 | -44.18 | 429.2832 | 3 | 3 | 7 | 1.4799 |
| AK-778/43206297 | 70.5 | -5.413 | -43.67 | 386.4545 | 1 | 4 | 7 | 3.6064 |
| AN-652/43265108 | 58.56 | -4.855 | -43.12 | 291.7367 | 2 | 3 | 4 | 3.3312 |
| AM-879/41890730 | 110.76 | -4.948 | -42.77 | 354.4338 | 2 | 4 | 10 | 3.5844 |
| AQ-390/42708956 | 141.65 | -5.097 | -42.53 | 398.4653 | 3 | 5 | 5 | 2.6944 |
| AO-854/43461140 | 49.33 | -5.301 | -42.37 | 275.7373 | 2 | 2 | 4 | 2.7094 |
| AN-329/41290657 | 113.93 | -4.997 | -42.24 | 275.3343 | 4 | 2 | 5 | 2.0303 |
| AN-329/43450029 | 76.66 | -4.732 | -42.05 | 328.3711 | 2 | 4 | 8 | 1.5272 |
| AE-848/32324022 | 132.8 | -4.716 | -41.89 | 410.4304 | 2 | 2 | 9 | 0.8458 |
| AQ-390/42708944 | 141.65 | -4.949 | -41.8 | 348.4047 | 3 | 5 | 5 | 1.786 |
| AK-968/15605224 | 130.17 | -5.11 | -41.61 | 464.5212 | 1 | 4 | 7 | 2.5912 |
| AP-970/40910914 | 67.43 | -4.776 | -41.31 | 326.3988 | 2 | 3 | 7 | 3.1797 |
| AM-807/42946882 | 156.62 | -4.845 | -41.22 | 426.4713 | 1 | 3 | 7 | 2.8767 |
| AN-329/43448916 | 70.23 | -4.846 | -41.21 | 331.8049 | 3 | 2 | 5 | 4.0665 |
| AG-670/11900377 | 95.44 | -5.399 | -41.11 | 345.3984 | 1 | 3 | 7 | 3.2499 |

| | | | | | | | | |
|---|---|---|---|---|---|---|---|---|
| AE-848/15341142 | 100.79 | -5.161 | -40.81 | 428.4282 | 0 | 5 | 8 | 3.4417 |
| AE-411/41415582 | 92.86 | -5.218 | -40.77 | 351.4274 | 2 | 3 | 5 | 4.007 |
| AN-652/41714081 | 49.33 | -5.029 | -40.48 | 255.3194 | 2 | 2 | 4 | 3.3008 |
| AO-548/43242488 | 49.33 | -4.98 | -40.34 | 269.3465 | 2 | 2 | 4 | 3.787 |
| AG-205/32365064 | 70.56 | -4.791 | -39.84 | 311.387 | 2 | 3 | 6 | 2.5081 |
| AK-968/15360853 | 120.94 | -5.078 | -39.57 | 418.4676 | 1 | 3 | 7 | 3.3256 |
| AN-465/40854064 | 62.83 | -4.75 | -39.39 | 349.2222 | 2 | 3 | 6 | 3.7354 |
| AK-968/15359278 | 120.94 | -5.14 | -39.37 | 386.4501 | 1 | 3 | 6 | 2.7713 |
| AS-871/40197285 | 50.36 | -4.858 | -39.02 | 242.2799 | 2 | 2 | 5 | 3.0426 |
| AO-548/43242520 | 58.56 | -4.882 | -38.93 | 271.3188 | 2 | 3 | 4 | 3.153 |
| AN-652/43024857 | 49.33 | -4.72 | -38.88 | 269.3465 | 2 | 2 | 4 | 3.787 |
| AO-365/43401572 | 119.33 | -4.786 | -38.82 | 278.335 | 2 | 4 | 6 | 0.6915 |
| AG-205/08215030 | 70.14 | -4.715 | -38.63 | 246.2461 | 2 | 3 | 3 | 1.5734 |
| AM-807/42946881 | 156.62 | -5.443 | -38.6 | 408.4809 | 1 | 3 | 7 | 2.6712 |
| AK-968/15254335 | 109.25 | -5.251 | -38.54 | 420.4489 | 2 | 4 | 8 | 2.6302 |
| AP-263/40045477 | 120.94 | -5.624 | -38.46 | 394.4294 | 1 | 3 | 7 | 3.1031 |
| AN-652/12058732 | 49.33 | -5.078 | -38.38 | 227.2652 | 2 | 2 | 3 | 2.3821 |
| AG-690/11450083 | 52.82 | -4.769 | -38.05 | 242.3003 | 3 | 2 | 3 | 1.5937 |
| AN-329/43385735 | 58.56 | -5.194 | -37.81 | 223.2742 | 2 | 3 | 3 | 3.2161 |
| AK-968/15604545 | 93.45 | -5.003 | -37.81 | 340.3619 | 1 | 3 | 5 | -0.1162 |
| AE-641/01074053 | 56.15 | -5.081 | -37.79 | 276.7901 | 2 | 0 | 5 | 4.7602 |
| AO-548/43179635 | 49.33 | -5.054 | -37.74 | 241.2923 | 2 | 2 | 3 | 2.8683 |
| AN-967/15488318 | 91.7 | -4.954 | -37.71 | 287.3398 | 2 | 4 | 5 | 2.6126 |
| AN-329/41402658 | 124.6 | -4.934 | -37.42 | 394.8382 | 3 | 3 | 7 | 0.8155 |
| AN-652/43340028 | 49.33 | -4.727 | -37.22 | 255.3194 | 2 | 2 | 3 | 3.3545 |
| AN-329/41402798 | 100.43 | -4.81 | -37.21 | 338.4315 | 2 | 2 | 4 | 3.1367 |

| | | | | | | | | |
|---|---|---|---|---|---|---|---|---|
| AK-918/40894305 | 95.5 | -4.775 | -37.18 | 367.4285 | 2 | 2 | 8 | 2.762 |
| AP-263/43479821 | 101.08 | -4.927 | -37.01 | 362.4278 | 1 | 3 | 6 | 3.0244 |
| AI-204/31695060 | 54.02 | -4.852 | -36.54 | 261.7131 | 2 | 2 | 4 | 2.7105 |
| AO-854/43464107 | 62.12 | -4.746 | -36.22 | 270.2655 | 1 | 3 | 5 | 2.5344 |
| AI-204/31695050 | 54.02 | -4.967 | -36.01 | 261.7131 | 2 | 2 | 4 | 2.9256 |
| AK-968/40708777 | 110.15 | -4.851 | -35.88 | 276.2538 | 2 | 3 | 5 | 1.4433 |
| AE-848/41826904 | 50.36 | -5.022 | -35.86 | 290.752 | 2 | 2 | 5 | 3.8447 |
| AI-204/31695051 | 54.02 | -4.695 | -35.48 | 261.7131 | 2 | 2 | 4 | 2.9256 |
| AI-204/31695049 | 54.02 | -5.018 | -35.46 | 245.2585 | 2 | 2 | 4 | 2.4667 |
| AO-080/43378936 | 84.22 | -4.894 | -34.95 | 378.4549 | 1 | 2 | 8 | 2.5876 |
| AE-641/01074059 | 69.04 | -4.702 | -34.84 | 243.3327 | 2 | 1 | 5 | 3.487 |
| AI-204/43372101 | 69.81 | -4.994 | -34.79 | 334.3034 | 3 | 2 | 5 | 3.5143 |
| AE-848/33212059 | 56.15 | -4.807 | -34.73 | 242.3451 | 2 | 0 | 5 | 4.0958 |
| AE-641/02615028 | 84.22 | -4.7 | -34.62 | 255.2786 | 3 | 2 | 4 | 2.4385 |
| AG-205/05140045 | 32.26 | -4.911 | -34.54 | 233.6997 | 2 | 1 | 3 | 3.6192 |
| AK-968/41017535 | 84.39 | -4.747 | -34.47 | 248.3708 | 2 | 0 | 5 | 3.6083 |
| AG-670/36669027 | 72.19 | -4.797 | -34.15 | 304.3516 | 2 | 2 | 4 | 2.5287 |
| AN-329/43385617 | 95.5 | -4.919 | -33.96 | 339.3743 | 2 | 2 | 8 | 1.3403 |
| AI-204/31695058 | 54.02 | -4.872 | -33.93 | 245.2585 | 2 | 2 | 4 | 2.2516 |
| AJ-292/41686516 | 79.27 | -4.716 | -33.76 | 293.3309 | 2 | 4 | 2 | 3.2451 |
| AG-690/12245523 | 137.73 | -5.32 | -33.6 | 345.4 | 2 | 2 | 7 | 1.3704 |
| AI-204/31696048 | 70.23 | -4.706 | -33.46 | 251.6743 | 3 | 2 | 3 | 2.1668 |
| AG-667/37281053 | 123.74 | -5.459 | -33.09 | 419.9104 | 2 | 2 | 6 | 3.3573 |
| AO-548/43196591 | 49.33 | -4.827 | -33.06 | 257.3353 | 2 | 2 | 5 | 2.3431 |
| AR-434/42808033 | 81.73 | -4.713 | -33.03 | 303.3607 | 2 | 2 | 3 | 3.0967 |
| AG-205/14151056 | 161.93 | -6.24 | -32.64 | 476.5568 | 2 | 3 | 8 | 3.3067 |

| | | | | | | | | |
|---|---|---|---|---|---|---|---|---|
| AN-329/41473712 | 112.16 | -4.792 | -32.43 | 375.4265 | 1 | 3 | 8 | 0.793 |
| AN-465/12944087 | 106.44 | -5.748 | -32.04 | 251.3092 | 2 | 3 | 5 | 0.8728 |
| AG-690/11669105 | 150.05 | -5.23 | -31.99 | 341.387 | 1 | 4 | 6 | 0.655 |
| AN-329/40278680 | 99.24 | -4.885 | -31.64 | 271.3432 | 3 | 1 | 5 | 3.3376 |
| AN-329/40396890 | 56.15 | -5.131 | -31.61 | 274.3626 | 2 | 0 | 5 | 4.6788 |
| AN-988/14610037 | 100.02 | -5.417 | -31.51 | 326.3348 | 2 | 3 | 7 | 2.5066 |
| AN-329/11638032 | 95.5 | -4.75 | -31.31 | 311.3201 | 2 | 2 | 6 | 0.8678 |
| AG-667/37281063 | 123.74 | -5.382 | -31.27 | 421.4988 | 2 | 2 | 6 | 3.1451 |
| AE-848/32733037 | 55.45 | -4.73 | -31.12 | 230.3124 | 3 | 1 | 2 | 1.9509 |
| AK-968/15612267 | 107.53 | -5.5 | -31.09 | 297.7421 | 1 | 2 | 5 | 1.7934 |
| AO-080/43441866 | 105.17 | -5.181 | -31.09 | 321.3125 | 1 | 4 | 6 | 0.5856 |
| AP-263/40971777 | 112.93 | -4.706 | -31 | 375.3409 | 1 | 2 | 6 | 2.0281 |
| AN-329/41435025 | 70.56 | -5.225 | -30.93 | 422.2896 | 2 | 3 | 3 | 4.0697 |
| AK-968/40344462 | 69.29 | -4.749 | -30.83 | 250.2967 | 2 | 1 | 5 | 3.2556 |
| AO-080/13769438 | 40.71 | -4.712 | -30.82 | 237.3069 | 1 | 1 | 4 | 2.3265 |
| AS-662/43412815 | 89.95 | -5.148 | -30.74 | 330.3667 | 1 | 2 | 6 | 0.7417 |
| AE-848/36287024 | 49.94 | -4.692 | -30.66 | 253.3063 | 2 | 2 | 4 | 3.4362 |
| AN-329/43449393 | 72.19 | -5.021 | -30.61 | 268.3181 | 2 | 2 | 4 | 2.1065 |
| AM-807/42946590 | 119.59 | -4.922 | -30.52 | 460.5109 | 0 | 3 | 6 | 3.9531 |
| AN-329/43448266 | 104.73 | -5.38 | -30.45 | 367.3848 | 2 | 3 | 6 | 1.1082 |
| AP-906/42698391 | 103.29 | -4.824 | -29.98 | 320.3465 | 1 | 3 | 5 | 1.1897 |
| AP-263/43503308 | 110.31 | -5.446 | -29.93 | 370.791 | 1 | 4 | 7 | 1.7276 |
| AO-022/43454301 | 97.11 | -5.263 | -29.88 | 353.4042 | 1 | 3 | 7 | 2.2563 |
| AK-968/41026299 | 123.74 | -4.733 | -29.79 | 321.3777 | 2 | 2 | 4 | 0.7031 |
| AO-095/42800782 | 40.71 | -4.794 | -29.56 | 257.7248 | 2 | 1 | 2 | 4.2544 |
| AK-968/13031191 | 107.53 | -5.122 | -29.53 | 277.3242 | 1 | 2 | 5 | 1.6152 |

| | | | | | | | | |
|---|---|---|---|---|---|---|---|---|
| AE-848/34780028 | 81.31 | -4.798 | -29.37 | 242.3043 | 2 | 2 | 3 | 3.2941 |
| AJ-292/15586002 | 95.94 | -5.28 | -29.19 | 319.3402 | 1 | 3 | 6 | -0.3496 |
| AK-968/12116299 | 118.43 | -4.899 | -28.92 | 291.3513 | 1 | 2 | 4 | 1.7871 |
| AG-205/08946044 | 95.5 | -5.107 | -28.87 | 311.3201 | 2 | 2 | 6 | 1.2892 |
| AO-365/09869013 | 62.22 | -4.995 | -28.74 | 228.2528 | 2 | 3 | 3 | 1.3885 |
| AS-871/42969578 | 91.29 | -5.179 | -28.41 | 423.3256 | 0 | 5 | 8 | 3.9464 |
| AK-968/11987351 | 70.08 | -4.695 | -27.95 | 355.4173 | 0 | 2 | 4 | 1.0318 |
| AN-329/14296008 | 104.73 | -5.126 | -27.93 | 391.1442 | 2 | 3 | 6 | -0.0888 |
| AM-807/43276841 | 147.83 | -4.764 | -27.71 | 466.5366 | 0 | 3 | 6 | 3.679 |
| AE-848/31937062 | 112.08 | -5.378 | -26.84 | 292.2924 | 2 | 3 | 4 | 0.9769 |
| AO-022/43453495 | 97.11 | -4.771 | -26.75 | 353.4042 | 1 | 3 | 7 | 2.2563 |
| AG-690/11426179 | 60.94 | -4.729 | -26.73 | 225.2521 | 3 | 2 | 2 | 2.8364 |
| AG-690/37071327 | 91.28 | -5.562 | -26.4 | 241.6774 | 1 | 1 | 3 | 1.5382 |
| AQ-390/43490286 | 112.08 | -5.185 | -26.06 | 356.38 | 2 | 3 | 4 | 2.3715 |
| AQ-086/41476304 | 78.01 | -4.91 | -25.96 | 252.2547 | 2 | 1 | 3 | 2.0304 |
| AQ-360/12120585 | 60.85 | -5.315 | -25.88 | 295.7482 | 0 | 1 | 4 | 1.0156 |
| AQ-360/40024608 | 60.85 | -4.964 | -25.59 | 261.3032 | 0 | 1 | 4 | 0.3512 |
| AK-918/40894318 | 95.5 | -4.824 | -25.37 | 339.3743 | 2 | 2 | 6 | 1.8496 |
| AK-968/41169476 | 56.15 | -4.72 | -25.13 | 331.3248 | 1 | 2 | 4 | 4.3533 |
| AO-990/37423036 | 73.74 | -5.048 | -24.96 | 328.3258 | 0 | 2 | 3 | 1.2834 |
| AS-768/43420458 | 88.76 | -4.813 | -24.7 | 303.3176 | 0 | 1 | 7 | 1.4017 |
| AP-434/40950083 | 92.86 | -4.965 | -24.37 | 299.309 | 1 | 2 | 7 | 2.1216 |
| AT-057/43485277 | 87.07 | -4.791 | -24.26 | 300.2937 | 1 | 3 | 3 | -0.4334 |
| AE-641/00652060 | 128.44 | -4.789 | -24.1 | 341.451 | 0 | 2 | 10 | 1.4074 |
| AE-848/04958048 | 89.95 | -4.782 | -23.93 | 350.3571 | 1 | 2 | 4 | 1.4825 |
| AG-690/12134089 | 137.73 | -4.766 | -23.65 | 343.3841 | 2 | 2 | 4 | 1.8716 |

| | | | | | | | | |
|---|---|---|---|---|---|---|---|---|
| AP-263/41686904 | 103.7 | -4.873 | -23.03 | 311.2969 | 1 | 1 | 6 | 0.796 |
| AG-690/15434541 | 150.98 | -5.746 | -22.42 | 362.429 | 0 | 2 | 9 | 2.1976 |
| AO-567/41089997 | 123.93 | -4.854 | -22.17 | 337.719 | 2 | 2 | 4 | 0.0066 |
| AJ-292/36376019 | 151.87 | -5.119 | -16.86 | 324.4028 | 1 | 3 | 3 | 1.4578 |
| AE-562/40806920 | 132.13 | -4.719 | -7.1 | 189.1026 | 1 | 1 | 5 | -3.4284 |

**Supplemental Table 2:** Screening Results of the VP40 Protein Based on Docking

| Name | TPSA | docking score | MMGBSA dG Bind | Molecular weight | Hydrogen bond donors | Hydrogen bond acceptors | Number of rotatable bonds | AlogP |
|---|---|---|---|---|---|---|---|---|
| AN-465/43369160 | 54.63 | -5.499 | -44.93 | 296.777 | 2 | 3 | 6 | 1.5243 |
| AN-465/43411361 | 59.95 | -5.443 | -43.47 | 330.407 | 2 | 4 | 7 | 1.5153 |

| | | | | | | | | |
|---|---|---|---|---|---|---|---|---|
| AN-465/43369952 | 45.4 | -5.932 | -43.15 | 301.195 | 2 | 2 | 5 | 2.2051 |
| AN-465/42888489 | 93.81 | -5.466 | -43.05 | 331.395 | 3 | 4 | 9 | -0.0005 |
| AN-465/43411466 | 40.44 | -6.457 | -42.87 | 356.298 | 2 | 1 | 6 | 0.6964 |
| AN-465/43411465 | 65.63 | -5.881 | -42.64 | 331.222 | 3 | 3 | 6 | 1.5219 |
| AN-465/42886175 | 42.96 | -6.635 | -42.49 | 269.367 | 0 | 3 | 8 | -0.1035 |
| AN-465/42886412 | 33.73 | -5.848 | -41.54 | 251.352 | 0 | 2 | 7 | 0.4114 |
| AN-465/43369750 | 45.4 | -6.568 | -40.25 | 287.168 | 2 | 2 | 5 | 1.8276 |
| AN-465/43369378 | 45.4 | -6.064 | -39.84 | 301.195 | 2 | 2 | 5 | 2.2051 |
| AN-465/43369098 | 45.4 | -5.466 | -39.77 | 315.222 | 2 | 2 | 6 | 2.7288 |
| AN-465/42246655 | 41.49 | -6.241 | -39.23 | 341.26 | 2 | 2 | 7 | 2.871 |
| AN-465/42886727 | 33.73 | -6.114 | -38.09 | 223.297 | 0 | 2 | 5 | -0.2862 |
| AN-465/42889349 | 85.61 | -6.728 | -38.04 | 317.819 | 3 | 3 | 10 | -1.9744 |
| AN-465/43384196 | 93.71 | -5.629 | -38.04 | 426.345 | 2 | 3 | 7 | 2.9455 |
| AN-465/42837116 | 76.82 | -6.43 | -37.94 | 376.302 | 2 | 3 | 10 | -1.6845 |
| AN-465/43411600 | 45.4 | -5.479 | -37.49 | 342.849 | 2 | 2 | 6 | 3.2616 |
| AN-465/43369242 | 57.43 | -5.538 | -37.38 | 344.264 | 2 | 2 | 8 | 1.8079 |
| AH-262/36336008 | 52.57 | -6.019 | -37.2 | 361.895 | 2 | 2 | 8 | 2.5503 |
| AN-465/42890043 | 93.81 | -5.547 | -37.12 | 365.84 | 3 | 4 | 9 | 0.6639 |
| AN-465/42833654 | 80.57 | -5.416 | -36.9 | 317.433 | 2 | 2 | 7 | 1.1833 |
| AO-365/43402886 | 74.58 | -5.622 | -36.88 | 447.345 | 2 | 3 | 7 | 2.8442 |
| AN-465/42886931 | 15.27 | -5.752 | -36.77 | 193.315 | 0 | 0 | 5 | 0.4319 |
| AN-465/43411631 | 93.71 | -5.625 | -36.71 | 426.345 | 2 | 3 | 7 | 2.9455 |
| AN-465/43369648 | 28.41 | -6.019 | -36.58 | 353.311 | 1 | 1 | 7 | 0.9132 |
| AN-465/43369437 | 45.4 | -5.401 | -36.45 | 284.741 | 2 | 2 | 5 | 1.7462 |
| AO-365/43474069 | 82.59 | -5.507 | -36.32 | 337.405 | 3 | 3 | 7 | 0.4127 |
| AN-465/42889971 | 42.52 | -5.563 | -35.65 | 368.883 | 2 | 2 | 10 | 0.9921 |

| | | | | | | | | |
|---|---|---|---|---|---|---|---|---|
| AK-778/11862141 | 32.34 | -5.787 | -35.62 | 262.161 | 1 | 1 | 4 | 0.9449 |
| AN-465/43369130 | 45.4 | -5.504 | -35.61 | 250.295 | 2 | 2 | 5 | 1.0818 |
| AN-465/42885888 | 55.41 | -5.537 | -35.46 | 351.88 | 2 | 3 | 10 | -0.1521 |
| AG-690/40081994 | 101.55 | -5.801 | -35.02 | 269.283 | 1 | 0 | 7 | 0.0679 |
| AS-813/43501852 | 71.55 | -6.46 | -35 | 209.191 | 2 | 1 | 2 | 3.4941 |
| AG-690/33049056 | 52.82 | -5.671 | -34.89 | 264.307 | 3 | 2 | 2 | 2.1177 |
| AO-365/43403046 | 41.57 | -6.339 | -34.82 | 233.293 | 1 | 2 | 4 | 0.3982 |
| AN-465/43461093 | 67.52 | -5.61 | -34.6 | 339.418 | 2 | 4 | 8 | 1.8346 |
| AN-465/42246715 | 82.81 | -5.877 | -34.54 | 329.807 | 2 | 4 | 8 | -0.0501 |
| AN-465/43369252 | 45.4 | -5.861 | -34.45 | 286.397 | 2 | 2 | 5 | 2.2133 |
| AN-465/43369210 | 45.4 | -5.65 | -34.41 | 300.425 | 2 | 2 | 5 | 2.6695 |
| AN-465/42887164 | 85.61 | -6.453 | -34.35 | 362.275 | 3 | 3 | 10 | -1.8904 |
| AN-465/43384165 | 33.29 | -5.409 | -34.11 | 330.449 | 2 | 1 | 7 | 0.8319 |
| AG-912/02706027 | 32.26 | -5.942 | -33.96 | 194.299 | 2 | 1 | 5 | 0.5013 |
| AP-970/42897159 | 89.37 | -6.225 | -33.87 | 388.878 | 3 | 3 | 9 | 1.2735 |
| AN-465/43411477 | 102.94 | -6.053 | -33.69 | 387.481 | 2 | 4 | 8 | 1.6003 |
| AN-465/43384150 | 42.52 | -6.101 | -33.6 | 360.476 | 2 | 2 | 8 | 0.8155 |
| AO-365/11349014 | 58.56 | -5.937 | -33.11 | 356.489 | 2 | 3 | 10 | 2.5015 |
| AN-465/43369233 | 45.4 | -5.465 | -33.09 | 325.233 | 2 | 2 | 6 | 2.1484 |
| AN-465/43369387 | 28.41 | -5.492 | -32.85 | 312.818 | 1 | 1 | 7 | 0.5485 |
| AN-465/43411506 | 48.64 | -6.615 | -32.79 | 304.436 | 2 | 2 | 8 | -0.5484 |
| AN-465/43384110 | 63.86 | -5.516 | -32.73 | 352.414 | 2 | 4 | 6 | 2.1628 |
| AN-465/43384109 | 73.64 | -5.884 | -32.41 | 314.43 | 2 | 2 | 6 | 1.9072 |
| AN-465/42886465 | 15.27 | -5.888 | -32.04 | 248.178 | 0 | 0 | 5 | 1.2745 |
| AE-641/00777058 | 42.85 | -6.272 | -31.83 | 240.264 | 1 | 3 | 3 | 3.0521 |
| AN-465/43384126 | 34.4 | -5.467 | -31.64 | 326.342 | 1 | 2 | 6 | 2.3393 |

| | | | | | | | | |
|---|---|---|---|---|---|---|---|---|
| AN-465/43370002 | 60.5 | -6.216 | -31.62 | 186.298 | 2 | 1 | 5 | -0.2038 |
| AN-465/43411181 | 63.22 | -5.88 | -31.55 | 326.422 | 2 | 3 | 8 | 0.433 |
| AS-813/43501794 | 78.34 | -7.047 | -31.54 | 184.172 | 2 | 2 | 2 | 0.3977 |
| AN-465/43369551 | 58.29 | -5.571 | -31.45 | 309.391 | 2 | 3 | 6 | 1.4592 |
| AO-080/42479511 | 110.02 | -5.713 | -31.43 | 231.272 | 2 | 4 | 4 | 1.1624 |
| AQ-239/43399392 | 75.21 | -5.814 | -31.34 | 218.214 | 2 | 4 | 3 | 1.7668 |
| AN-465/43411505 | 139.44 | -5.416 | -31.29 | 372.407 | 4 | 6 | 8 | -0.304 |
| AO-476/15577245 | 141 | -5.906 | -31.13 | 439.044 | 1 | 2 | 6 | 3.7282 |
| AG-777/36176008 | 55.48 | -5.58 | -30.62 | 218.278 | 2 | 2 | 4 | 0.3775 |
| AI-204/31677002 | 56.73 | -6.752 | -30.32 | 245.734 | 2 | 2 | 6 | -0.3425 |
| AN-465/43369277 | 93.71 | -5.507 | -30.32 | 357.455 | 2 | 3 | 7 | 1.6167 |
| AG-670/13120074 | 32.34 | -5.592 | -30.23 | 211.261 | 1 | 1 | 4 | -0.1784 |
| AN-465/43421754 | 82.7 | -8.241 | -30.12 | 371.824 | 2 | 2 | 7 | 1.702 |
| AN-465/43384151 | 42.52 | -5.656 | -30.04 | 360.476 | 2 | 2 | 8 | 0.8155 |
| AN-465/43461029 | 84.58 | -5.568 | -29.95 | 303.384 | 3 | 3 | 8 | 0.2133 |
| AJ-292/41083288 | 91.41 | -5.917 | -29.84 | 358.489 | 1 | 4 | 4 | 1.2918 |
| AE-641/01943051 | 114.79 | -6.188 | -29.79 | 188.206 | 3 | 3 | 1 | -0.5102 |
| AQ-086/43478989 | 96.58 | -5.676 | -29.32 | 207.256 | 3 | 1 | 1 | 1.5746 |
| AL-182/11270030 | 126.63 | -6.313 | -28.57 | 250.239 | 0 | 2 | 3 | -1.026 |
| AI-204/31709041 | 37.28 | -5.438 | -28.34 | 215.232 | 1 | 2 | 3 | 2.8807 |
| AN-465/43421554 | 93.81 | -5.429 | -28.28 | 361.465 | 3 | 4 | 7 | -0.2703 |
| AI-204/33264023 | 71.3 | -5.379 | -27.8 | 211.246 | 1 | 1 | 5 | -0.4977 |
| AO-365/43403048 | 41.57 | -7.187 | -27.68 | 259.331 | 1 | 2 | 4 | 0.8592 |
| AN-465/43384111 | 45.4 | -5.775 | -27.31 | 326.394 | 2 | 2 | 6 | 2.6002 |
| AN-465/42887163 | 76.82 | -5.753 | -27.11 | 361.267 | 1 | 3 | 9 | -0.5162 |
| AK-823/41252445 | 55.48 | -5.522 | -26.95 | 196.272 | 2 | 2 | 4 | 0.3052 |

| | | | | | | | | |
|---|---|---|---|---|---|---|---|---|
| AB-323/13887426 | 118.56 | -5.835 | -26.91 | 210.239 | 3 | 3 | 1 | 0.4263 |
| AN-465/43369998 | 93.71 | -5.406 | -26.5 | 436.356 | 2 | 3 | 7 | 2.3651 |
| AI-204/43489630 | 108.2 | -6.107 | -26.32 | 265.362 | 2 | 3 | 5 | -0.0263 |
| AE-641/00358058 | 71.55 | -5.812 | -26.22 | 232.327 | 2 | 1 | 3 | 1.7821 |
| AN-465/43411025 | 82.7 | -8.24 | -26.02 | 337.379 | 2 | 2 | 7 | 1.0376 |
| AN-465/43421952 | 33.29 | -5.419 | -25.85 | 270.805 | 2 | 1 | 5 | -0.2927 |
| AE-641/01923022 | 94.56 | -6.051 | -24.76 | 200.261 | 2 | 2 | 2 | 0.6996 |
| AN-465/43421896 | 77.66 | -5.4 | -24.48 | 319.427 | 3 | 3 | 9 | 0.2521 |
| AN-465/43411020 | 90.71 | -6.711 | -24.37 | 313.378 | 1 | 1 | 6 | 1.0608 |
| AN-465/43421895 | 82.7 | -8.474 | -24.36 | 351.406 | 2 | 2 | 7 | 1.3213 |
| AN-465/42246534 | 93.81 | -5.907 | -23.79 | 303.768 | 3 | 4 | 8 | -0.5708 |
| AN-465/43411021 | 80.93 | -5.741 | -23.52 | 381.432 | 1 | 3 | 9 | 1.8367 |
| AN-465/43411023 | 80.93 | -7.706 | -23.45 | 351.362 | 1 | 3 | 6 | 1.3164 |
| AP-263/43505353 | 57.79 | -5.93 | -23.33 | 305.849 | 1 | 2 | 6 | 1.0183 |
| AA-516/12432323 | 54.26 | -5.413 | -23.28 | 190.289 | 1 | 0 | 2 | 0.962 |
| AN-329/43449771 | 35.25 | -5.741 | -23.18 | 190.626 | 1 | 1 | 3 | 0.8497 |
| AN-465/42243976 | 50.72 | -5.405 | -22.15 | 224.282 | 2 | 3 | 5 | 0.2669 |
| AK-968/11533152 | 73.04 | -5.425 | -22.13 | 252.278 | 3 | 2 | 3 | 1.8909 |
| AG-690/10108038 | 6.48 | -5.394 | -22.13 | 322.498 | 0 | 0 | 6 | 1.871 |
| AN-465/42519030 | 50.72 | -6.534 | -21.59 | 274.77 | 2 | 3 | 7 | 1.1304 |
| AQ-390/43364049 | 86.89 | -6.272 | -21.54 | 362.474 | 2 | 3 | 8 | 0.0696 |
| AI-346/33252018 | 47.25 | -6.387 | -21.48 | 270.334 | 0 | 1 | 4 | -0.4972 |
| AE-641/01119004 | 63.37 | -5.522 | -21.18 | 193.21 | 1 | 2 | 1 | 1.7935 |
| AN-465/43421641 | 62.47 | -7.494 | -19.19 | 325.343 | 1 | 1 | 6 | 1.7538 |
| AH-357/04334061 | 26.02 | -5.561 | -18.7 | 161.128 | 1 | 0 | 1 | 2.0257 |
| AJ-797/43492786 | 81.49 | -5.387 | -18.23 | 264.305 | 1 | 2 | 3 | -1.0369 |

| | | | | | | | | |
|---|---|---|---|---|---|---|---|---|
| AN-465/43421587 | 75.36 | -7.478 | -18.13 | 308.34 | 1 | 2 | 6 | 0.3977 |
| AP-123/42300618 | 44.01 | -6.225 | -18.13 | 199.211 | 0 | 1 | 2 | -0.1781 |
| AN-465/43384225 | 62.47 | -5.777 | -17.89 | 325.343 | 1 | 1 | 6 | 1.7538 |
| AL-182/11270028 | 83.49 | -5.742 | -17.82 | 219.268 | 0 | 2 | 2 | -0.4342 |
| AN-465/43411022 | 62.47 | -7.885 | -17.61 | 231.253 | 1 | 1 | 4 | -0.0352 |
| AO-365/41690590 | 32.34 | -5.389 | -16.47 | 298.21 | 1 | 1 | 4 | 0.8255 |
| AO-365/43473996 | 96.25 | -5.53 | -16.3 | 222.291 | 2 | 2 | 3 | -0.3504 |
| AI-204/43372130 | 28.16 | -6.279 | -16.25 | 251.762 | 2 | 0 | 4 | 0.9638 |
| AG-690/40698438 | 140.28 | -5.804 | -15.54 | 446.919 | 2 | 3 | 9 | 3.6128 |
| AN-465/42886076 | 15.27 | -6.026 | -15.53 | 275.34 | 0 | 0 | 8 | 1.5856 |
| AN-465/43411016 | 62.47 | -7.008 | -14.83 | 307.352 | 1 | 1 | 6 | 1.5483 |
| AE-641/00356033 | 110.11 | -6.227 | -14.65 | 356.467 | 2 | 4 | 7 | -0.7469 |
| AO-022/43453140 | 103.88 | -5.704 | -14.43 | 457.622 | 1 | 5 | 6 | 2.0651 |
| AN-465/43411011 | 62.47 | -6.947 | -14.37 | 287.362 | 1 | 1 | 7 | 1.5453 |
| AP-124/43383762 | 64.07 | -5.693 | -13.48 | 192.243 | 2 | 2 | 3 | -0.8383 |
| AN-465/43411240 | 65.12 | -6.121 | -13.34 | 294.356 | 2 | 0 | 6 | 0.746 |
| AP-782/41885444 | 135.34 | -5.696 | -12.93 | 216.111 | 0 | 2 | 3 | 0.2739 |
| AK-968/15254087 | 60.49 | -6.316 | -12.81 | 285.372 | 2 | 2 | 3 | -0.3957 |
| AH-357/02166042 | 58.88 | -5.884 | -11.2 | 202.258 | 2 | 1 | 4 | 1.5257 |
| AG-690/11482436 | 56.06 | -6.551 | -9.21 | 298.388 | 2 | 1 | 5 | 1.8775 |
| AS-871/43475746 | 78.76 | -5.751 | -8.55 | 315.334 | 0 | 6 | 3 | 0.2867 |
| AN-329/43333566 | 51.29 | -5.855 | -6.09 | 260.362 | 3 | 1 | 4 | -0.5763 |
| AE-641/30177043 | 15.71 | -6.061 | -5.14 | 252.403 | 0 | 1 | 7 | -0.3968 |
| AO-476/43414255 | 65.01 | -6.139 | -0.83 | 314.39 | 1 | 2 | 4 | 0.4591 |
| AE-848/34417033 | 95.49 | -5.613 | 0.78 | 268.172 | 0 | 4 | 2 | 0.3719 |

**Supplemental Table 3:** Molecular docking of interacting amino acids between the chemicals and EBOV VP35/40 protein

| System | Hydrophobic interactions | Hydrogen bonds |
|---|---|---|
| 1ES6_AN-465_ 43369160 | ASP-144, PRO-196 | GLY-198,GLU-325 |
| 1ES6_AN-465_ 43369952 | ASP-144, PRO-196 | GLY-198,GLU-325 |
| 1ES6_AN-465_ 43411361 | ASP-310,PRO-196 | ASP-312,GLU-325,LYS-326 |
| 3FKE_AN-465_ 43369198 | ILE-295,PRO-293,LYS-248,VAL-245 | GLN-244,GLN-241 |
| 3FKE_AN-465_ 43369333 | ILE-295,PRO-304,LYS-248,PRO-293 | GLN-244,GLN-241 |
| 3FKE_AO-022_ 43452438 | PRO-293, PRO-304, ILE-295, GLN-244, LYS-248 | GLN-244,ARG-225,TYR-229,GLN-241 |
